# Clustering individuals using INMTD: a novel versatile multi-view embedding framework integrating omics and imaging data

**DOI:** 10.1101/2024.09.23.614478

**Authors:** Zuqi Li, Sam F. L. Windels, Noël Malod-Dognin, Seth M. Weinberg, Mary L. Marazita, Susan Walsh, Mark D. Shriver, David W. Fardo, Peter Claes, Nataša Pržulj, Kristel Van Steen

## Abstract

**Motivation:** Combining omics and images, can lead to a more comprehensive clustering of individuals than classic single-view approaches. Among the various approaches for multi-view clustering, nonnegative matrix tri-factorization (NMTF) and nonnegative Tucker decomposition (NTD) are advantageous in learning low-rank embeddings with promising interpretability. Besides, there is a need to handle unwanted drivers of clusterings (i.e. confounders).

**Results:** In this work, we introduce a novel multi-view clustering method based on NMTF and NTD, named INMTD, that integrates omics and 3D imaging data to derive unconfounded subgroups of individuals. In the application to real-life facial-genomic data, INMTD generated biologically relevant embeddings for individuals, genetics and facial morphology. By removing confounded embedding vectors, we derived an unconfounded clustering with better internal and external quality; the genetic and facial annotations of each derived subgroup highlighted distinctive characteristics. In conclusion, INMTD can effectively integrate omics data and 3D images for unconfounded clustering with biologically meaningful interpretation.

**Availability and implementation:** https://github.com/ZuqiLi/INMTD

## 1 Introduction

Clustering is a crucial technique in data analysis, enabling the identification of intrinsic structures within complex datasets by grouping similar data points. In the field of medicine, clustering has been widely used for uncovering disease subtypes, tailoring personalized treatments, and improving early diagnosis (Ghosal *et al*. 2020). As data complexity grows, clustering methods based on a single view or single data source are often insufficient, necessitating the development of more sophisticated approaches. Multi-view clustering has emerged as a powerful solution, leveraging multiple data perspectives to enhance clustering quality and reveal richer patterns than single-view methods (Rappoport and Shamir 2018; Chauvel *et al*. 2020). As the data views commonly used in biomedical science to describe an individual, omics and imaging data have shown essential advantages in understanding various biological phenomena (Antonelli *et al*. 2019). For instance, Chen et al. obtained better prediction for subtypes of lung adenocarcinoma by integrating extracted features from histopathological images and omics data than using a single view (Chen *et al*. 2021). However, few people have worked on clustering individuals based on omics and imaging data. Moreover, processing images normally requires tensor methods due to their 3D format (Hériché, Alexander and Ellenberg 2019), e.g. a color image consists of pixels represented by height, width and color channel, and a 3D mesh consists of X, Y, Z coordinates for height, weight and depth.

Various multi-view clustering methods have been developed, which can be generally classified into three categories, based on the relationship between data integration and clustering (Rappoport and Shamir 2018): 1) early integration combines all datasets into a single one before building the model for clustering, 2) intermediate integration clusters a joint embedding learnt from all views, and 3) late integration computes a clustering from each dataset and then merges all clusterings together. Out of the three, intermediate integration approaches have shown superior performance in many applications possibly because they require a model specifically designed for multi-view clustering tasks (Khan and Maji 2019; Wang *et al*. 2020; Yun *et al*. 2021). By clustering subjects with multi-view data from a joint embedding, integrative nonnegative matrix factorization (intNMF) proposed by Chalise and Fridley has found similar cancer subtypes identified by previous studies (Chalise and Fridley 2017). This embedding represents the subjects with patterns that are naturally additive and hence easily interpretable (Lee and Seung 1999). A well-established extension of NMF model for better interpretation and clustering is nonnegative matrix tri-factorization (NMTF) that decomposes the input dataset into three smaller matrices (Ding *et al*. 2006). However, NMTF models only work with 2D matrices and cannot deal with data views of higher dimensions, e.g. a 3D tensor, which is the common data format for imaging or spatial data. A generalization of NMTF to tensors is the nonnegative Tucker decomposition (NTD) (Kim and Choi 2007). It decomposes the original input tensor into a core tensor with the same number of dimensions and one embedding matrix corresponding to each of its dimensions. There have been attempts to integrate multiple cross-linked 2D matrices into a tensor, which, however, does not work on originally 3D data (Luo *et al*. 2022). Another work by Broadbent et al. combined a tensor with a similarity matrix, while they treated the matrix as a graph regularization to the Tucker decomposition instead of a separate data view (Broadbent, Song and Kuang 2024).

Clustering real-world data is often complicated by the presence of confounders—factors that influence the observed data in an unwanted way (Liu *et al*. 2015). Confounders can obscure the true clustering structure, leading to spurious results (Schwarz *et al*. 2024). Addressing confounders is essential for accurate clustering, as their effects can mask the genuine patterns within the data. Effective clustering methods must account for these confounding variables to uncover the true underlying structure. A widely adopted approach is to regress out the confounding effects from every feature during pre-processing, but it comes with the potential loss of useful signals prior to the modelling (Pourhoseingholi, Baghestani and Vahedi 2012). Some other strategies include kernel conditional clustering which computes the final clustering conditioned on confounders (He *et al*. 2020). However, the conditional clustering is computationally expensive and cannot efficiently work on high-dimensional data.

Here in this paper, we propose a novel multi-view clustering method, integrative non-negative matrix and tensor decomposition (INMTD), which obtains unconfounded clustering jointly from 2D and 3D datasets. It learns an embedding matrix for each data dimension and subgroups the individuals from their embedding after removing vectors in the embedding space that are linked with confounders. Because the true cluster structure of real-life patient dataset is often unknown, we evaluated INMTD on a US cohort from healthy individuals (White *et al*. 2021), whose heterogeneity mainly comes from the population structure with confounders including age, sex, etc. Combining 2D genotypes and 3D facial morphology, our model computed biologically meaningful embeddings and connected well the facial and genetic embeddings. Furthermore, INMTD derived an unconfounded clustering of individuals with better intrinsic quality and clearer association with population structure than the original clustering. We also characterized each population subgroup with their enriched genetic pathways and highlighted facial areas.

## 2 Methods

### 2.1 INMTD: integrative non-negative matrix and tensor decomposition with correction for confounders

INMTD unifies NMTF and NTD to cluster subjects with multi-view data of 2D and 3D structure. We assume *p*_1_ subjects described by two data views, a 2D matrix 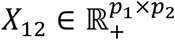 of *p*_2_ features and a 3D tensor 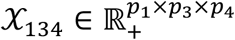 of *p*_3_ features in the 2nd dimension and *p*_4_ features in the 3rd dimension, both nonnegative. The aim of our method is to jointly compute the embedding matrices for each dimension and cluster the *p*_1_ subjects based on its own embedding (Fig. 1).

**Fig. 1:**
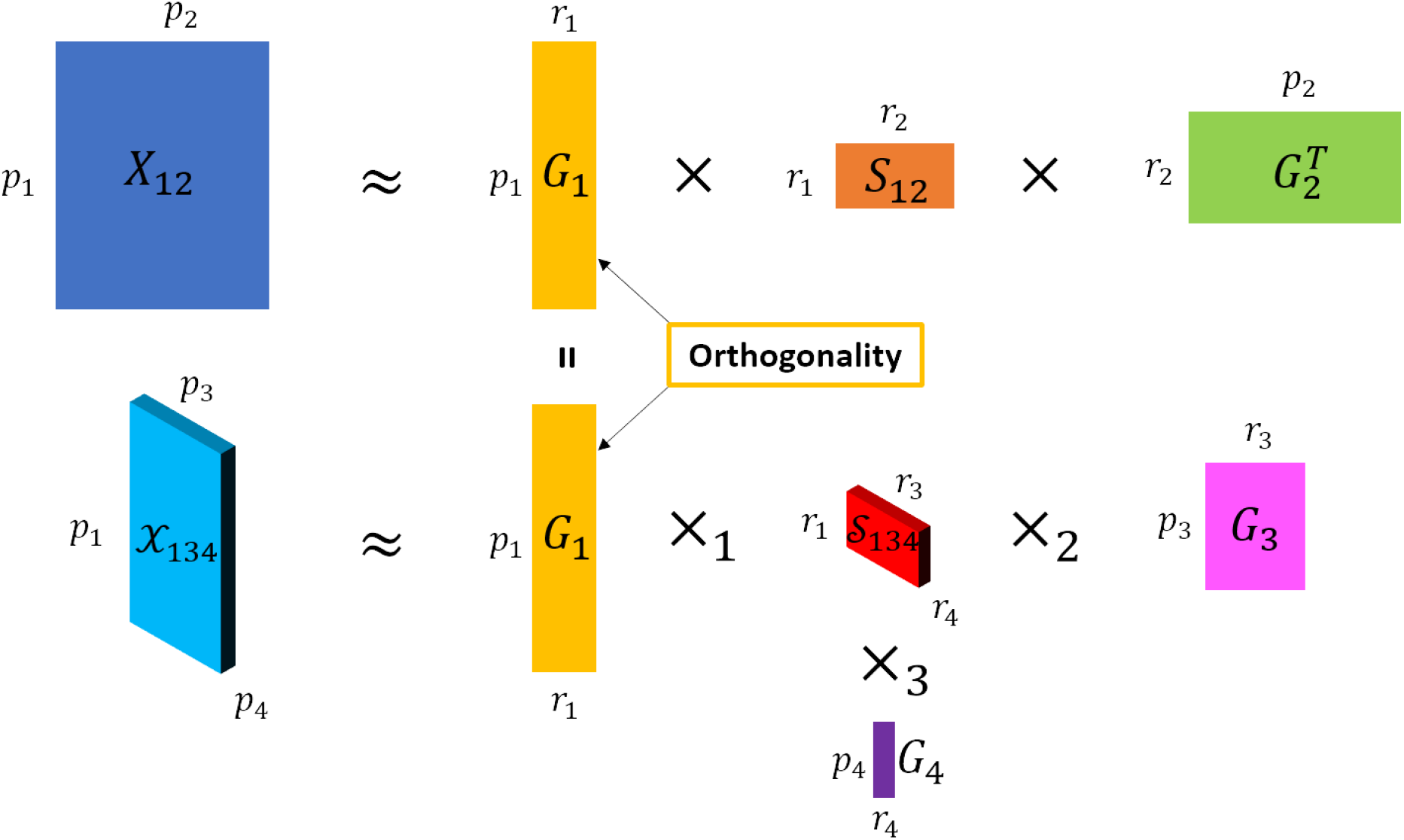
Overview of INMTD model for integrating 2D and 3D data. The 2D matrix *X*_12_ is decomposed into two embedding matrices *G*_1_ and *G*_2_ and a core matrix *S*_12_. The 3D tensor 𝒳_134_ is decomposed into three embedding matrices *G*_1_, *G*_3_ and *G*_4_ and a core tensor 𝒮_134_. INMTD integrates *X*_12_ and 𝒳_134_ by jointly optimizing *G*_1_, which is shared by the two views. The orthogonality constraint on *G*_1_ ensures disentanglement of embedding vectors.

For 2D matrix *X*_12_, NMTF factorizes it into three nonnegative submatrices 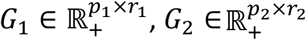 and 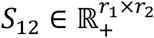, so that 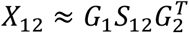. The objective of NMTF is to find the optimal *G*_1_, *G*_2_ and *S*_12_ that minimize the reconstruction error:

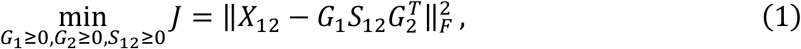

where 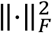 indicates the Frobenius norm and ≥ 0 for a matrix means all values in that matrix should be nonnegative. The multiplicative update rules to solve this objective function have been proposed by Ding et al. (Ding *et al*. 2006) *G*_1_ and *G*_2_ are low-rank embeddings for the *p*_1_ subjects and *p*_2_ features, respectively, where *r*_1_ and *r*_2_ are their ranks and normally *r*_1_ ≪ *p*_1_ and *r*_2_ ≪ *p*_2_. *S*_12_ is the core matrix that links *G*_1_ with *G*_2_ and can be considered as the compressed representation of *X*_12_.

Similar to NMTF, the NTD model decomposes the 3D tensor 𝒳_134_ into 3 embeddings 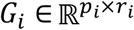 with *i* ∈ {1,3,4}, named the mode matrices, and a core tensor 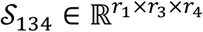 using the mode product of tensor:

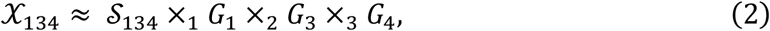

where 𝒮_134_ ×_*n*_ *G*_*i*_ is the mode-*n* product between tensor 𝒮_134_ and matrix *G*_*i*_, resulting in a new tensor with its *n* -th dimension changed. The objective of NTD is to minimize the reconstruction error:

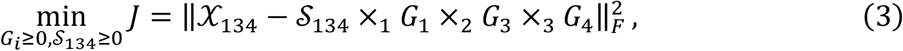

To jointly decompose *X*_12_ and 𝒳_134_, we derive an integrative objective from the two views via NMTF and NTD:

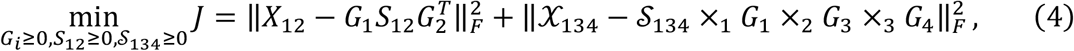

where 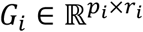 are the low-rank embeddings corresponding to *p*_1_ subjects, *p*_2_ features of view 1, *p*_3_ features of view 2 and *p*_4_ channels of view 2. The rank parameters *r*_*i*_ are determined via the rule of thumb: 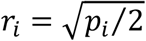 (Kodinariya and Makwana 2013). *G*_1_ is shared by both terms in formula (4) and jointly learnt from both views. We further adopt an orthogonality constraint on *G*_1_ for more rigorous clustering interpretation (Ding *et al*. 2006):

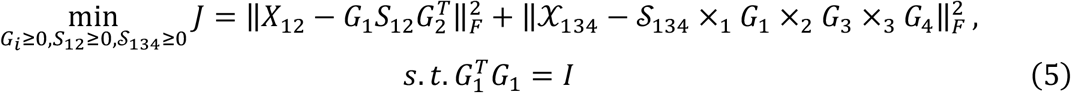

Because formula (5) has no analytic solution for *G*_*i*_, *S*_12_ and 𝒮_134_, we iteratively compute their values via the multiplicative update rules (Dissez *et al*. 2019) (see section 2.2). Furthermore, in each iteration, we normalize every column of *G*_1_ after updating to further guarantee unit vectors and eliminate the scale indeterminacy as suggested by Li et al. (Bo Li, Guoxu Zhou and Cichocki 2015)

We apply *k*-means clustering (with 10 random initializations) on the joint embedding, *G*_1_, and select the best number of clusters based on Silhouette score. Silhouette score is a classic internal metric that measures how well a dataset is clustered. A higher value (close to 1) suggests a more valid clustering.

To assess how much *G*_1_ is confounded by a set of known confounders, *C*, we conduct a linear model F-test between every confounder and every column of *G*_1_. The unconfounded clustering is then recomputed from the columns of *G*_1_ that have no significant association with any confounders. This is done also by *k*-means and Silhouette score for the optimal number of clusters. The removal of confounded embedding components is based on the additive nature of NMF-based methods that the data is represented by the sum of all its embedding aspects (Lee and Seung 1999). Deconfounding at the embedding level is computationally less expensive and can better preserve meaningful information than the widely used approach, which is to regress out confounders from each feature at the input level.

### 2.2 Training procedure of INMTD

To solve INMTD, we use a fixed-point method that, starting from an initial solution, iteratively uses multiplicative update rules to converge towards a locally optimal solution. During the optimization process, all the embedding matrices and core matrix/tensor of INMTD are iteratively updated to minimize the objective function (formula (5)). Following the derivation procedure used to derive multiplicative update rules for orthogonal NMTF and NTD, we derive the update rules for INMTD:

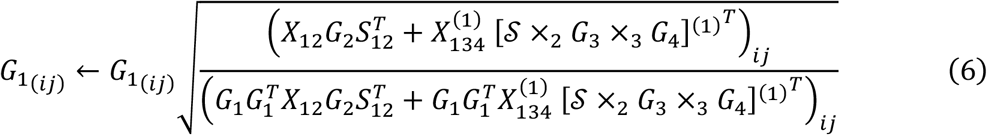

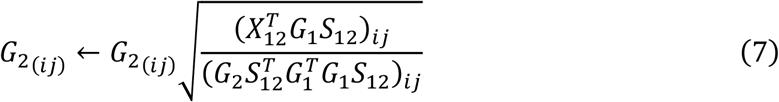

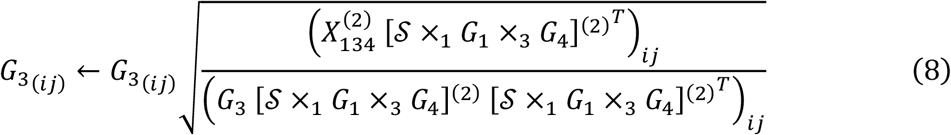

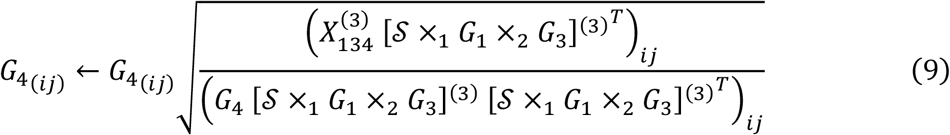

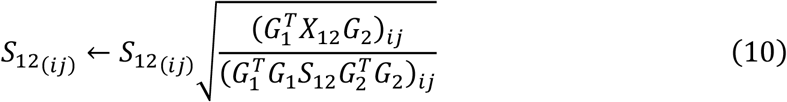

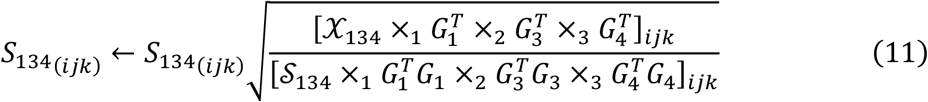

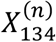 denotes the mode-*n* matricization of 𝒳_134_, which reshapes 𝒳_134_ to a 2D matrix along its *n*-th dimension.

We initialize *G*_*i*_ via singular value decomposition (SVD), which has shown better results than random initialization (Malod-Dognin *et al*. 2019). For *X*_12_, the original matrix is decomposed by SVD and *G*_1_ and *G*_2_ are derived from the left and right matrices of SVD, respectively. Because 𝒳_134_ is 3D, we run SVD for every slice along the *p*_4_ channels and average all the right matrices to compute the initial *G*_3_ and similarly run SVD for every slice along the *p*_3_ features to initialize *G*_4_. As the update rules are multiplicative, to avoid entries in *G*_*i*_ to remain 0, we add an infinitesimal number (1e-5) so that these entries can be updated.

*S*_12_ and 𝒮_134_ can also be initialized by SVD through the eigen values if the original data frames are symmetric. But we don’t assume their symmetry in our framework, therefore, we apply the following rules:

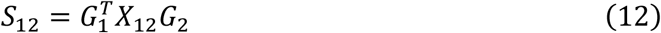

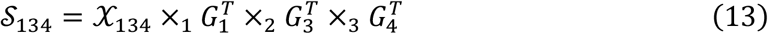

The initialized *S*_12_ and 𝒮_134_ are automatically nonnegative because all the multipliers are nonnegative.

To assess the goodness and convergence of INMTD, we track a metric along the optimization, which is the total relative error:

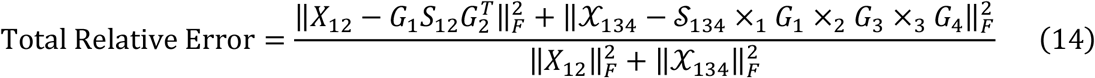

The total relative error computes the fraction of the reconstruction errors of the two datasets *X*_12_ and 𝒳_134_ in their L2 norms. It is a nonnegative value and a lower total relative error indicates better reconstruction from the decomposed elements.

### 2.3 Association between embeddings

Linking two embeddings from different views can be achieved by mapping them to the same space of *G*_1_ because *G*_1_ is the embedding shared by both data types. More specifically, we project *G*_2_ to the space of *G*_1_ via the core matrix *S*_12_, so that 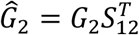. The new matrix Ĝ_2_now has the same embedding size as *G*_1_ and is in the same embedding space as *G*_1_. Similarly, *G*_3_ is also projected to the space of *G*_1_ via the core tensor 𝒮_134_, resulting in Ĝ_3_ = 𝒮_134_ ×_3_ *G*_3_. In the special case when *r*_4_ = 1 and hence 𝒮_134_ is of shape *r*_1_ × *r*_3_ × 1, this is equivalent to 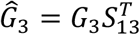, where *S*_13_ reshapes 𝒮_134_ to *r*_1_ × *r*_3_. Subsequently, the relationship between feature (row) *i* in *G*_2_ and feature (row) *j* in *G*_3_ can be assessed by cosine similarity:

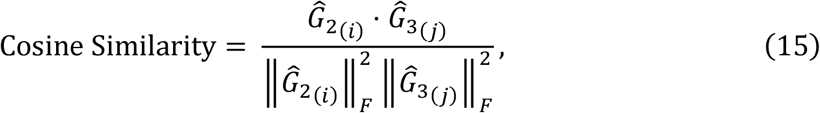

where 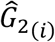 indicates the *i* -th row of Ĝ_2_ and 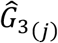 the *j* -th row of Ĝ_3_. Cosine similarity measures the dot product between two vectors regardless of their magnitudes, providing good normalization when comparing between Ĝ_2_ and Ĝ_3_. It ranges from -1 to 1 and the higher the more similar between the two vectors.

## 3 Experiments

### 3.1 Evaluation dataset

We apply INMTD to a multi-view dataset of 4,680 normal people with European ancestry, characterized by a 2D matrix and a 3D tensor (White *et al*. 2021). These people were recruited from three independent studies in the US, 3D Facial Norms cohort (PITT), Pennsylvania State University (PSU) and Indiana University-Purdue University Indianapolis (IUPUI). Every individual was described by 7,141,882 SNPs and a 3D mesh image which contains the X, Y, Z coordinates of 7,160 landmarks, namely *X*_12_ ∈ ℝ^4,680×7,141,882^ and 𝒳_134_ ∈ ℝ^4,680×7,160×3^, respectively. Due to the enormous number of SNPs the SVD initialization on *X*_12_ had to adopt randomized SVD for feasibility. The initialization on 𝒳_134_ still used full SVD. To standardize genomic and facial data, we subtracted the mean from each view and divided every entry by the maximum of each view. We then took the absolute values to ensure nonnegativity. The rank *r*_1_ of embedding *G*_1_ was determined via the rule of thumb: 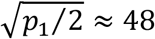 where *p*_1_ is the number of individuals, thus 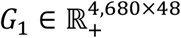. *r*_2_ was chosen in a similar way but based on the number of protein-coding genes, namely 19,430, instead of SNPs to reduce the computational burden, and thus 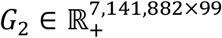. We had 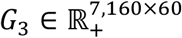 and 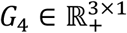 because 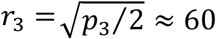 and 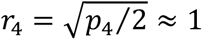 where *p*_3_ = 7,160 and *p*_4_ = 3. For every individual, we also collected a few covariates, including age, sex, height, weight, face size and camera system. BMI was derived via: BMI = weight(*kg*)/height(*m*)^2^ as an additional covariate. Here, we consider these covariates as confounders to population structure because they might hinder us in finding the population subgroups based on genetic heterogeneity. To measure the population structure of the cohort, White et al. have computed four ancestry axes by projecting the genomic data onto the principal component (PC) space of the SNPs from the 1000G Project (White *et al*. 2021).

### 3.2 Evaluation on the Embeddings

INMTD learns an embedding for SNPs, faces, and individuals. Here, we outline how we assess the biological validity for those embeddings separately and jointly.

To biologically validate our individual embedding, we assess if *G*_1_ captures any heterogeneity of the cohort, including both population structure (ancestry axes) and confounding effects. Due to the orthogonality constraint, embedding vectors of *G*_1_ can be considered as independent and characterize different aspects of the individuals. We, therefore, test the statistical association between every embedding vector and every ancestry axis or confounder. In particular, Kruskal-Wallis ANOVA is used for ancestry axes and continuous confounders (age, height, weight, BMI and face size) and chi-squared test is used for categorical confounders (sex and camera system). We apply Benjamini-Hochberg (BH) correction for the multiple testing.

To assess our SNP embedding, we cluster SNPs based on their embedding and use enrichment analysis to see if their space is functionally organised. As *G*_2_ is not orthogonal, we use *k*-means (with 10 random initializations) on *G*_2_ to subgroup SNPs into *r*_2_ = 99 clusters. We first check how well those SNP clusters coincide with the 19,430 protein-coding genes given the fact that gene is a natural summary of SNPs and biological processes are usually interpreted on a gene level. A gene is defined to be enriched in a SNP cluster if SNPs located 2K base pairs around this gene are present in this cluster significantly more than in the background. Hypergeometric test is applied for the enrichment analysis and the BH procedure is used for multiple testing correction. To further check if these clusters have biological relevance, we run another enrichment analysis for Gene Ontology (GO) terms for biological process. We annotate every SNP by GO terms if it is mapped to a gene that is annotated by a GO term and then test the overrepresentation of GO terms in every SNP cluster. Because GO terms are gene annotations, we annotate SNPs with GO terms that annotate their mapped genes. Only GO terms under biological process category are used and the BH procedure is applied for multiple testing correction.

To assess our facial embedding, facial landmarks are subgrouped by Ward’s hierarchical clustering on *G*_3_ into *r*_3_ = 60 clusters, which segments the shape of face. The Ward’s method has been shown outperforming other common linkage methods (Ferreira and Hitchcock 2009; Vijaya, Sharma and Batra 2019). We first compute the Ward distance between every pair of landmarks in *G*_3_, based on which we then construct a hierarchical tree and cut it at a height with 60 clusters. We adopt hierarchical clustering in order to compare with the hierarchical segmentation done on the same dataset by White et al. They segmented the facial shape from global to local into five levels with 63 segments. To compare the ability of the hierarchical tree of *G*_3_ and the hierarchical segmentation by White et al. to faithfully capture the pairwise dendrogrammatic distances between landmarks, we compute their cophenetic correlation coefficient:

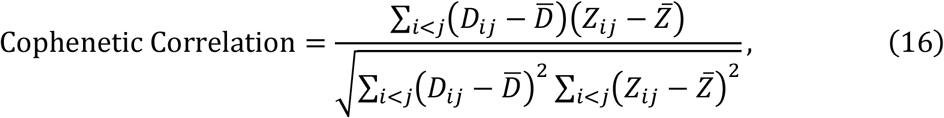

where *i* and *j* are facial landmarks, *D* is the Euclidean distance matrix between landmarks, and Z is the cophenetic distance matrix between landmarks, denoting the heights at which two points are first merged in the dendrogram. 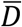 and 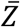 are the mean of *D* and Z, respectively. A cophenetic correlation close to 1 indicates a high-quality hierarchical clustering.

To assess the association between the SNP and facial embeddings, we map them to the space of *G*_1_ and compute cosine similarity between each SNP and each facial landmark (see section 2.3). For the SNPs closest to facial landmarks in the joint space, we apply the GREAT analysis (v.4.0.4), which finds biological meaning of the set of SNPs via the annotations of nearby genes. Technically speaking, GREAT analysis performs a binomial test for a set of SNPs to check whether the overlap between their associated genes and genes with a certain annotation is greater than random chance. To associate SNPs with genes, we apply the default and recommended settings (McLean *et al*. 2010), namely the ‘basal plus extension’ rule with 5kb upstream and 1kb downstream plus 1000kb extension. Note that one SNP can be associated with multiple genes and SNPs not associated with any genes are not included in this analysis.

### 3.3 Characterization of unconfounded population subgroups

The unconfounded population subgroups are characterized based on the projection of genetic and facial embeddings to the space of the sample embedding, enabling computing the similarity between population subgroup centroids and SNPs and facial landmarks. For each subgroup, we first select the top 0.1% SNPs with highest cosine similarity to its centroid in the joint space, to which the GREAT analysis is applied to reveal the most relevant phenotypes (HPO). The threshold of 0.1% is determined by balancing the number of genes selected per subgroup and the genomic coverage of genes selected for all subgroups (Supp. Fig. 1 and 2). We then visualize the cosine similarities of all facial landmarks to a subgroup centroid on the averaged face, in order to demonstrate how different areas of the face are associated with the corresponding subgroup.

## 4 Results

### 4.1 INMTD generates biologically meaningful embeddings from a real-life multiview dataset

We applied INTMD to a real-world facial-genomic cohort collected from the US for unconfounded population subgrouping (White *et al*. 2021). This dataset consists of two data types, 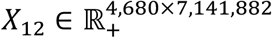 for 7,141,882 SNPs in 4,680 people and 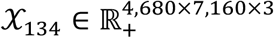 for the 3D coordinates of 7,160 facial landmarks in the same 4,680 individuals (White *et al*. 2021). A few confounders were collected as well, which are age, sex, height, weight, camera system and face size. We also derived BMI (body mass index) from height and weight as a potential confounder. To assess the population structure of this cohort, White et al. computed four ancestry axes by projecting the genotypes to a principal component space built from the 1000 Genomes Project data, in the manner of EIGENSTRAT (Price *et al*. 2006). We ran our INMTD model on this multi-view dataset for 1,000 iterations with SVD initialization and it converged in terms of the total relative error (Supp. Fig. 3). Due to the heuristic solver for formula (5), *G*_1_ is only approximately orthonormal. Therefore, we further checked the independencies between the embedding vectors of *G*_1_ based on its covariance matrix, namely 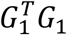 (Supp. Fig. 4).

Because the heterogeneity of a population can be largely described by population structure, age, sex, etc., we validated the information captured by *G*_1_ based on the statistical association between every column vector of *G*_1_ and every ancestry axis or confounder (Fig. 2). The results indicate that most vectors captured the information of population structure while being confounded. This is expected as, for instance, height has been reported to highly relate with different European ancestries (Cavelaars *et al*. 2000). Furthermore, the 48 embedding vectors have different association patterns with the ancestry axes and confounders. For instance, the 1st vector is significantly associated with most ancestry axes as well as confounders, while no association with the 6th vector is observed.

**Fig. 2:**
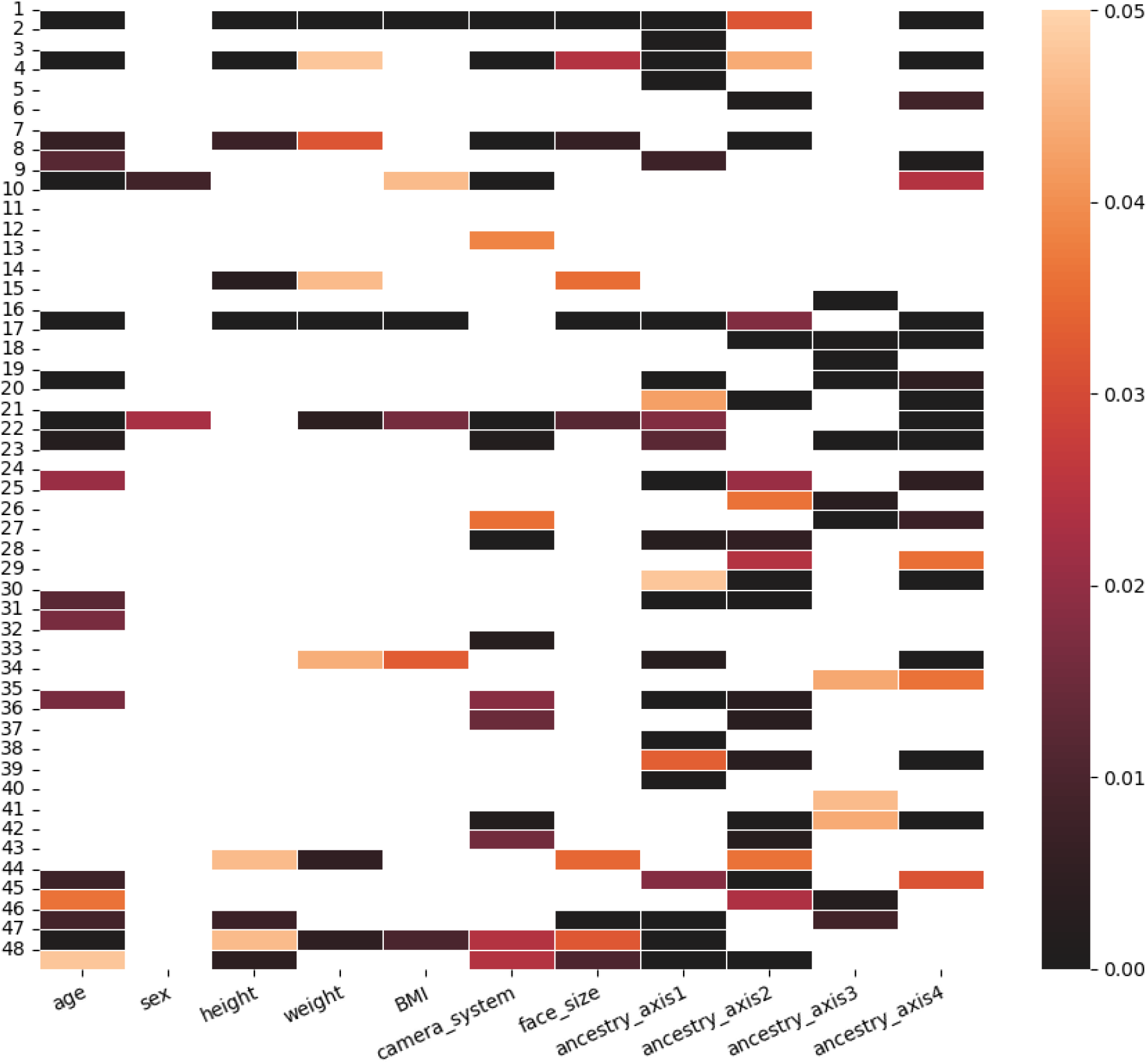
Heatmap of P-values for the linear F test between every *G*_1_ embedding vector and every confounder and ancestry axis. Non-significant P-values (larger than 0.05 after BH correction) were removed from the plot.

To assess the biological relevance of the SNP embedding from *G*_2_, we applied *k*-means on *G*_2_ to derive 99 clusters of SNPs. We first mapped every SNP to genes if it falls within or 2k base pairs around a gene and ignored SNPs that do not map to any genes. 97 out of 99 SNP clusters have at least one overrepresented gene (P-value < 0.05 in a hypergeometric test with BH correction for multiple testing) with respect to the background of all the SNPs that can map to a gene. 98.6% of genes have been enriched in at least one SNP cluster while most genes were enriched in only a few clusters (Fig. 3). We then applied GO (gene ontology) enrichment analysis for each cluster after assigning GO annotations of a gene to all its belonging SNPs. 96.0% of *G*_2_ clusters statistically significantly overrepresented at least one GO term and 99.6% of GO terms have been enriched in at least one cluster (Fig. 4). This result validated that the SNP clusters obtained from *G*_2_ have both genomic specificity and biological relevance.

**Fig. 3:**
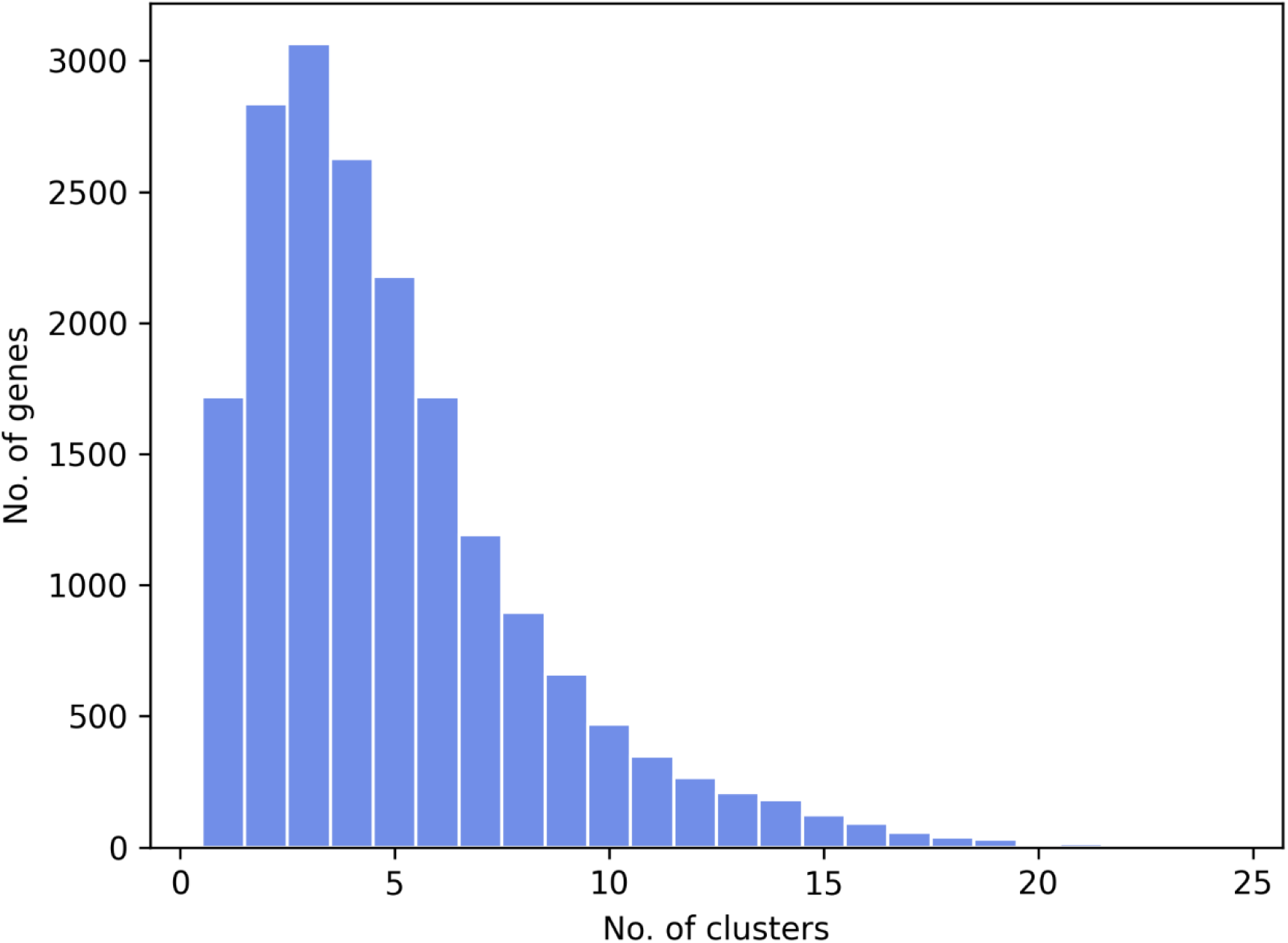
Histogram showing the number of genes (y-axis) that are enriched in a given number of *G*_2_ clusters (x-axis). An enrichment analysis is done between every gene and every *G*_2_ (SNP) cluster. We then compute how many genes (y-axis) are found to be enriched in different No. of clusters of SNPs (x-axis).

**Fig. 4:**
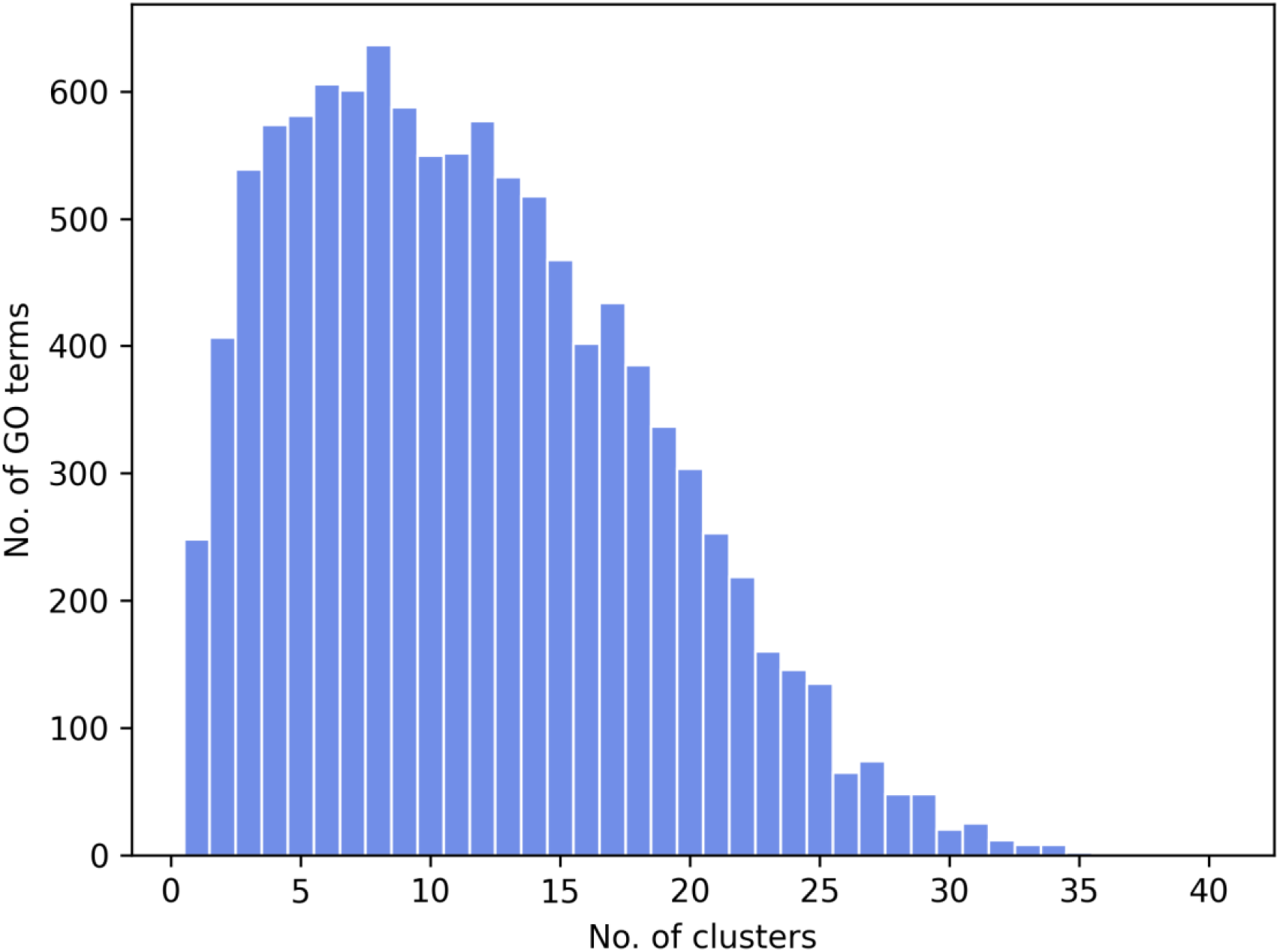
Histogram showing the number of GO terms (y-axis) that are enriched in a given number of *G*_2_ clusters. An enrichment analysis is done between every GO term and every *G*_2_ cluster. We then compute how many GO terms (y-axis) are found to be enriched in different No. of clusters of SNPs (x-axis).

To assess the quality of the facial embedding from *G*_3_, we aimed to obtain a hierarchical segmentation on *G*_3_ and compare it with the work of White et al. (White *et al*. 2021) The chosen Ward’s hierarchical clustering yielded 60 facial segments (Supp. Fig. 5), with a cophenetic correlation coefficient of 0.638, which is higher than that of the facial segmentation by White et al. on the same images (0.414). It suggested that the hierarchical segmentation from embedding *G*_3_ better groups together facial landmarks that are close in 3D space than the one by White et al., defining more spatially coherent ‘patches’. The visualization of these 60 clusters on an averaged face also showed that the landmarks within each cluster are spatially close to each other and the clustering is morphologically meaningful (Fig. 5).

**Fig. 5:**
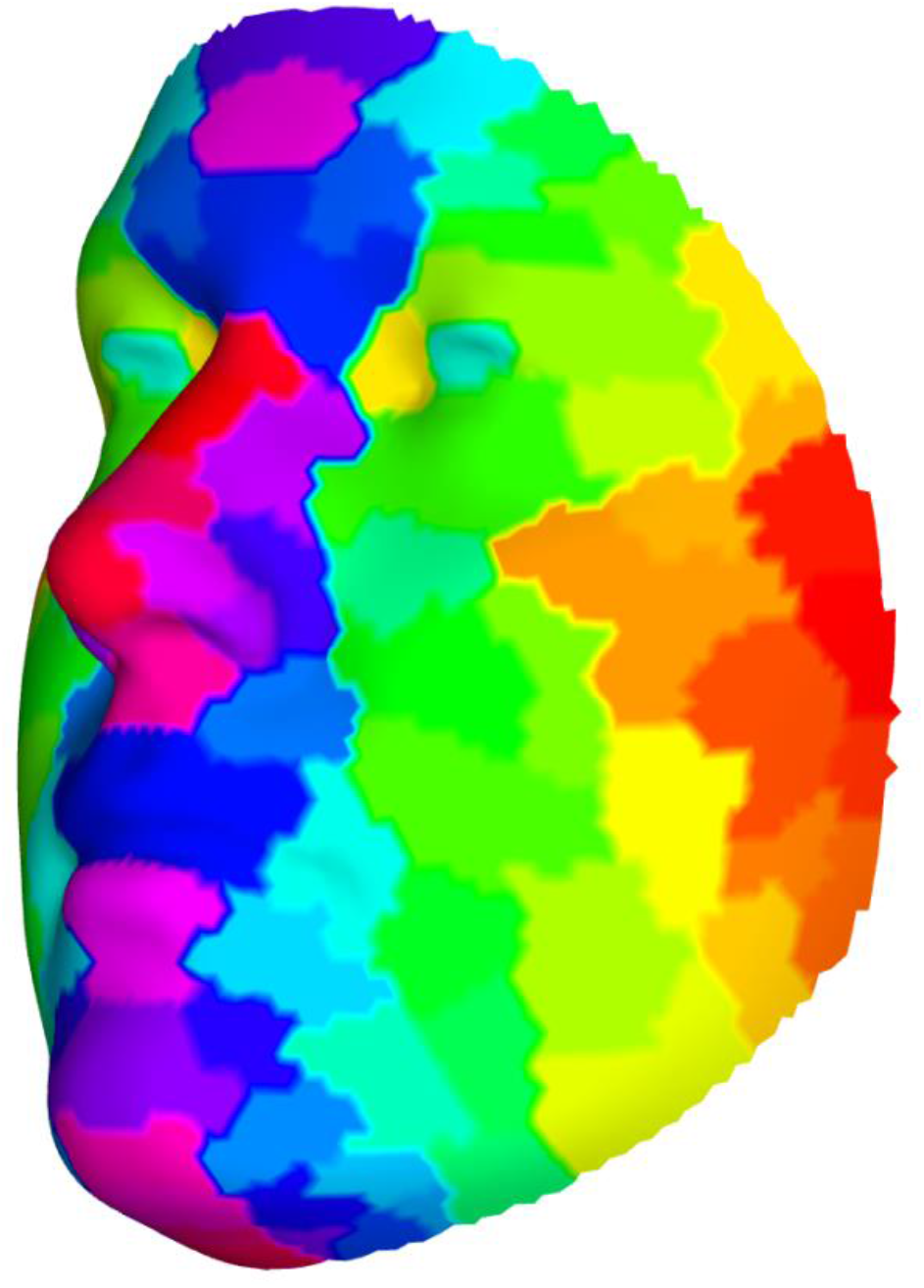
The 60 clusters derived from *G*_3_, illustrated on the mean face shape of all individuals. Not every cluster has a unique color due to the large number of clusters, but neighboring clusters are distinguished by different colors. Every face shape is symmetric along X-axis (left/right), so is the facial segmentation.

To further investigate the relationships between genetics and facial morphology, we mapped both *G*_2_ and *G*_3_ to the space of *G*_1_ (Supp. Fig. 6). For each facial landmark in the joint space, we selected the closest SNP in terms of cosine similarity, resulting in 905 unique SNPs in total. To assess what biological traits are associated with the chosen SNPs, we conducted a GREAT analysis on their neighbouring genes (McLean *et al*. 2010), which found 17 significantly enriched human phenotype ontology (HPO) terms (Gargano *et al*. 2024) based on the adjusted binomial P-values (Fig. 6). Most of the enriched terms are highly linked to facial morphology (especially eyes), limb or spine morphology and embryonic development, depicting close biological relationship between the embeddings for SNPs and facial landmarks, namely *G*_2_ and *G*_3_. Other enriched terms suggested high relatedness between facial morphology and myotonia. The results indicate that INMTD allows for uncovering biologically relevant associations between SNPs and facial landmarks.

**Fig. 6:**
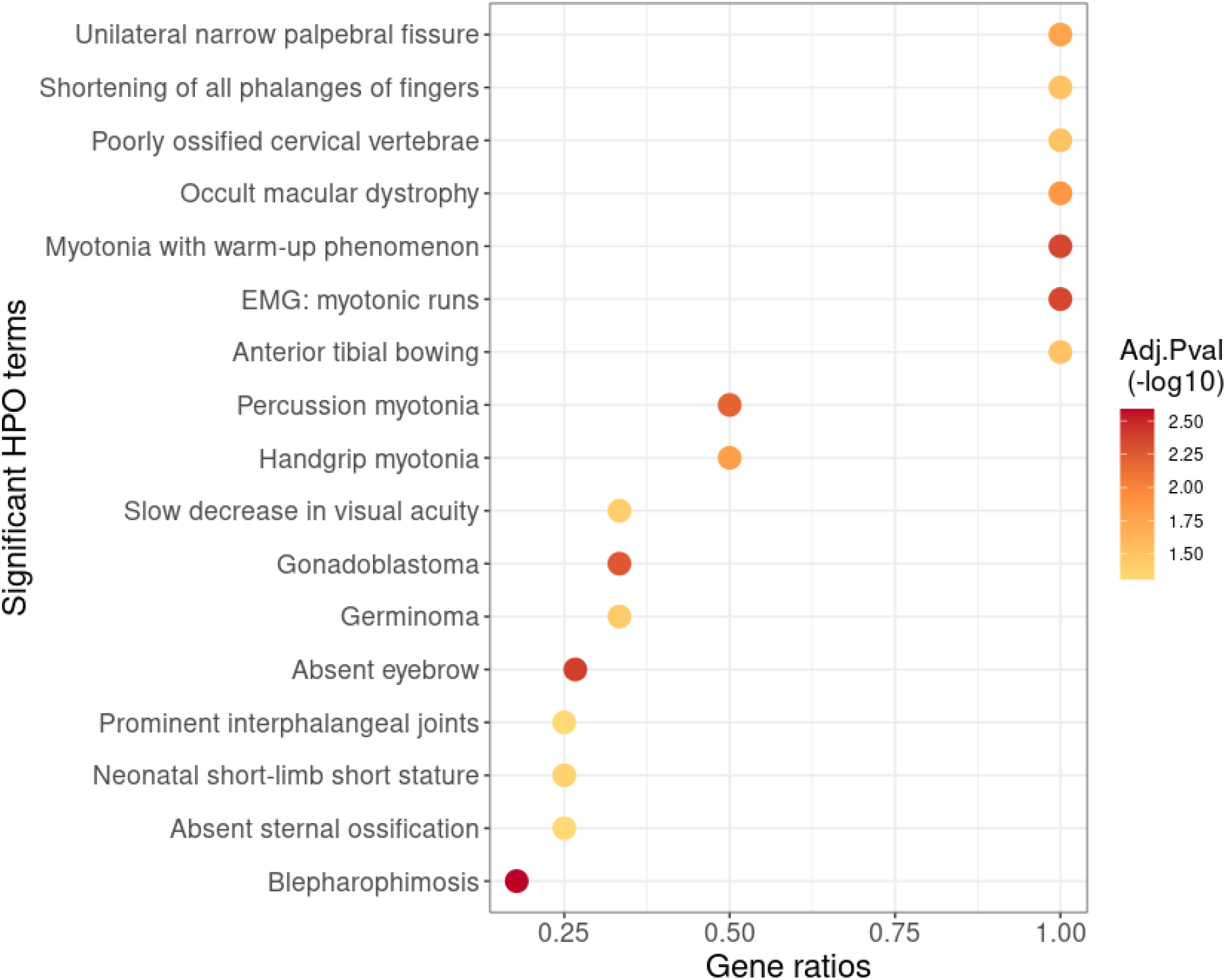
Dotplot of 17 significantly enriched HPO terms from 905 SNPs that are closest to facial landmarks in the joint space. GREAT analysis first mapped the 905 SNPs to 1,156 genes based on the default and recommended settings, and then ran a binomial test for the enrichment analysis with BH correction. X-axis shows the ratio between the number of observed genes and the number of genes annotated for each HPO term. The dot color indicates the significance of the adjusted binomial P-value in the form of -log10.

### 4.2 INMTD finds unconfounded population subgroups characterized by their genetic and facial information

To derive the optimal population subgroups, we applied *k* -means on *G*_1_ with different number of clusters and found that the clustering with 48 clusters achieved the highest Silhouette score (0.169) (Supp. Fig. 7). As mentioned before, many vectors of *G*_1_ are significantly confounded, potentially disturbing correct interpretation and characterization of derived subgroups. In order to deal with the confounding effect, we removed the 28 columns in *G*_1_ that are significantly associated with any confounders, and clustered individuals based on the remaining 20 embedding vectors (Supp. Fig. 8). We then applied *k*-means on the unconfounded *G*_1_ with different number of clusters and the clustering with 20 clusters achieved the highest Silhouette score (0.329) (Supp. Fig. 9). This score is statistically higher (empirical P-value = 0.02 from 1000 repetitions, Supp. Fig. 10) than clusterings with the same number of clusters from 20 randomly sampled *G*_1_ vectors, indicating better intrinsic validity of the unconfounded clustering.

To validate our reduced space is unconfounded and leads to a better capturing of the population structure, we assessed the statistical association between the derived clustering and the ancestry axes and confounders. The Kruskal-Wallis test (nonparametric ANOVA) showed both the original and the unconfounded clusterings are significantly associated with all ancestry axes (P-value< 0.05), suggesting their relationships with population structure (Table 1). Yet the original clustering also has significant associations with most confounders, especially age and camera system, while the unconfounded clustering has no significant associations with any confounders, validating our unconfounding strategy. The effective reduction of the influence by camera system, which resembles the batch effect, also indicates the strength of the confounder correction.

**Table 1:** Adjusted P-values of the statistical tests between the clustering of *G*_1_ and every ancestry axis and confounder. Kruskal-Wallis test was used for continuous variables and Chi-squared test for categorical variables. All P-values have been corrected for multiple testing via the Benjamini-Hochberg (BH) procedure. Column 3 are adjusted P-values from the original clustering based on all vectors in *G*_1_ while column 4 from the unconfounded clustering. Adjusted P-values lower than 0.05 (threshold for significance) are in bold.

| Variable | Statistical Test | Adj. P-value<br>(original) | Adj. P-value<br>(unconfounded) |
| --- | --- | --- | --- |
| Ancestry axis 1 | Kruskal-Wallis | **7.24e-40** | **2.99e-9** |
| Ancestry axis 2 | Kruskal-Wallis | **1.04e-11** | **2.29e-7** |
| Ancestry axis 3 | Kruskal-Wallis | **6.07e-11** | **2.69e-3** |
| Ancestry axis 4 | Kruskal-Wallis | **8.11e-22** | **8.16e-11** |
| Age | Kruskal-Wallis | **9.36e-8** | 0.888 |
| Sex | Chi-squared | 0.118 | 0.899 |
| Height | Kruskal-Wallis | 8.38e-2 | 0.888 |
| Weight | Kruskal-Wallis | 5.51e-2 | 0.951 |
| BMI | Kruskal-Wallis | **2.19e-3** | 0.899 |
| Camera system | Chi-squared | **3.01e-78** | 0.899 |
| Face size | Kruskal-Wallis | **3.72e-2** | 0.888 |

To investigate if the derived subgroups from the unconfounded *G*_1_ clustering capture well the population structure, we adopted 3,519 European ancestry informative markers (EuroAIMs) found by Tian et al. (Tian *et al*. 2009), which are SNPs capable of distinguishing European subpopulations. We first mapped the SNP embedding *G*_2_ to the space of *G*_1_ and then, in the joint space, selected 3,519 SNPs with highest cosine similarities to the centroids of derived subgroups, which drive the clustering of individuals. We selected the same number of SNPs as the EuroAIMs for better comparison. Because only 13 EuroAIMs were originally included in our dataset, we looked at the gene level and found that 856 out of the 3,519 selected SNPs are located in the same genes as the EuroAIMs. A hypergeometric test showed that this fraction is significantly (P-value = 2.15e-63) higher than a random selection from all SNPs in our dataset, indicating that the genetic basis of the derived subgrouping is statistically significantly associated with the European population structure. We also checked for the subgroups derived from the original *G*_1_ clustering and found only 40 out of 3,519 selected SNPs that are located in the EuroAIM genes. The P-value of 1 from the hypergeometric test also implied this fraction is not statistically higher than a random selection. This result further proved that our deconfounding strategy has successfully led to population subgroups that could better highlight the population structure.

After showing the validity of our unconfounded population subgrouping, we focused on the characterization of two subgroups that are most associated with the four ancestry axes (Supp. Table 1), namely subgroup 2 and 7. The top 0.1% SNPs selected for population subgroup 2 were significantly enriched in over 500 HPO terms, with the top 20 terms clearly related to skeletal morphology or bone formation (Fig. 7A). It is in line with the highlighted areas on the face (Fig. 7B), e.g., the nasal bone, the zygomatic bone, maxilla, and most part of the mandible. This result characterized population subgroup 2 with facial skeleton. Note that the frontal bone does not show up as much as the facial bones, implying the different morphology of cranial and facial skeleton (Anderson *et al*. 2024). Some other enriched HPO terms involve anemia, telangiectasia and neutrophil, which are related to the blood. Whereas the genetic representation of population subgroup 7 found 72 significantly associated HPO terms in total, and many of the top 20 terms are strongly involved in kidney function (Fig. 7C). Meanwhile, the eye area was remarkably underlined on the mean face (Fig. 7D), which supports the common embryogenic stage of eyes and kidneys and the reported relationship between eye and kidney diseases (Bodaghi, Massamba and Izzedine 2014). Therefore, population subgroup 7 is likely characterized by eye morphology related to kidney function. Other enriched terms indicate the involvement of kidney function in diabetes, hair development, tooth development, etc.

**Fig. 7:**
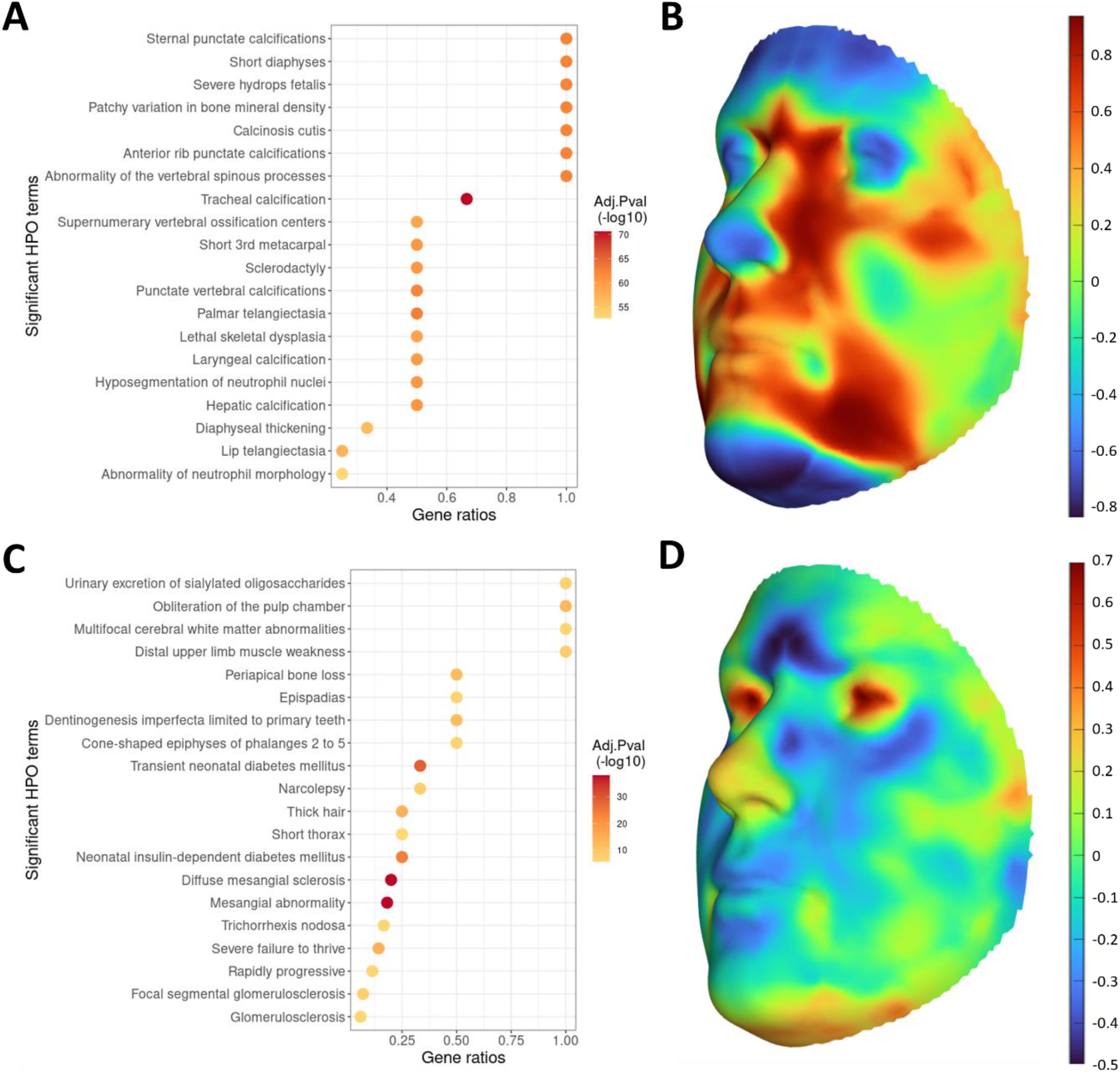
Genetic and facial representation for population subgroup 2 (top, A and B) and 7 (bottom, C and D). The genetic representation was obtained via GREAT analysis (left) for the top 20 HPO terms enriched in the top 0.1% SNPs highlighted for population subgroup 2 (A) and 7 (C). We used the default and recommended settings of GREAT analysis with binomial test and BH correction. For the facial representation, the cosine similarity between each facial landmark and the centroid of subgroup 2 (B) and 7 (D) separately was plotted on the mean face. Red color indicates higher value while blue color indicates lower value.

## 5 Discussion and conclusion

In this study, we proposed INMTD, a framework that integrates both 2D matrices and 3D tensors for unconfounded clustering, and applied it to a real-life facial-genomic dataset to evaluate its performance and find an unconfounded subgrouping for European population structure. We derived 20 unconfounded population subgroups with their representative genetic and facial characteristics, providing the potential for more precise healthcare towards each subpopulation.

This work applies INMTD to facial and genomic data, which reflect a large fraction of the population structure. A comprehensive subgrouping of population structure has the potential to facilitate precision medicine where individuals in each population subgroup may receive tailored medical decisions based on their intrinsic characteristics (Saria and Goldenberg 2015; Loh, Cao and Zhou 2019). For instance, special medical treatment may be given to individuals in subgroup 2 for their different skeleton development and individuals in subgroup 7 for their distinct kidney function.

While classic matrix or tensor decomposition models only focus on a single dataset and current integrative matrix factorization models cannot deal with higher-dimensional data structures, INMTD is able to jointly decompose both matrix and tensor data. Another key feature of INMTD is its orthogonality constraint on *G*_1_ for better clustering and interpretation. An orthogonal matrix has all its vectors independent to each other, and therefore, every vector can be investigated specifically for its characteristics. We further normalized the vectors of *G*_1_ to make them orthonormal, resembling naturally a cluster indicator matrix. Optionally, we can impose orthogonality on any embedding if needed.

As an extension of joint NMF model, INMTD has the potential to predict facial images from genotypes or vice versa of new samples, in a similar fashion as Akata et al. (Akata, Thurau and Bauckhage 2011) In the former case, given the genotypes 𝓍_12_ of an individual that has not been seen by the model, we first solve 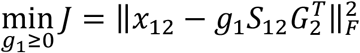 for *g*_1_ and then predict its facial image 𝓍_134_ = 𝒮_134_ ×_1_ *g*_1_ ×_2_ *G*_3_ ×_3_ *G*_4_. Alternatively, we could find the closest image from the cohort to 𝓍_134_ according to cosine similarity or the image whose corresponding embedding vector in *G*_1_ space is closest to *g*_1_. In addition, a new sample can be classified into one of the derived subgroups by assigning *g*_1_ to its closest cluster centroid.

Even though INMTD was illustrated in a facial-genomic dataset for population subgrouping, it is not restricted to facial images and genomic data. INMTD can be applied to any 2D and 3D datasets for joint clustering, as long as they are non-negative or can be converted to non-negative values, e.g. transcriptomics or epigenomics as 2D matrices and CT scans or time series data as 3D tensors. It is also possible to further extend INMTD to deal with more than two data types or data of higher dimensions, e.g. a moving 3D image (4D), by adding extra embeddings to the model and objective function.

There are two main limitations of INMTD for future improvements. The first one is that it determines the ranks of each embedding based on a rule of thumb because of the extremely large scale of genomic dataset. Nevertheless, in most cases the data dimensionality would be feasible for INMTD model on a modern computer or computing server and the user could choose the optimal ranks via cross-validation or other non-heuristic methods. The other potential deficiency comes from the post-hoc confounder correction, which removes confounded vectors but in a fairly strict manner with the risk of ‘over-correcting’. A more compromising strategy could be adding a regularization term in the objective function to iteratively minimize the confounding effect in *G*_1_, which indicates our future work.

In conclusion, with the surge in biological data in diverse formats and the growing demand for personalized medicine with Big Data, INMTD is envisaged to become an essential tool for integrating multi-view datasets of varying dimensions, enabling meaningful and unconfounded clustering. We are confident that INMTD has the potential of widespread adoption in the future due to its exceptional performance and ease-of-interpretation.

## Data availability

For the 3D Facial Norms dataset, genotypic markers are available to the research community through the dbGaP controlled-access repository (http://www.ncbi.nlm.nih.gov/gap) at accession #phs000929.v1.p1. The raw source data for the phenotypes - the 3D facial surface models in .obj format - are available through the FaceBase Consortium (https://www.facebase.org) at accession #FB00000491.01. Access to these 3D facial surface models requires proper institutional ethics approval and approval from the FaceBase data access committee. The PSU and IUPUI datasets were not collected with broad data sharing consent. Code for INMTD is publicly available on GitHub: https://github.com/ZuqiLi/INMTD.

## Acknowledgements

This project has received funding from the European Union’s Horizon 2020 research and innovation programme under the Marie Sklodowska-Curie grant agreements No. 813533 (MLFPM) and No 860895 (TranSYS). We also want to acknowledge the European Research Council (ERC) Consolidator Grant No. 770827, the Spanish State Research Agency and the Ministry of Science and Innovation MCIN grant PID2022-141920NB-I00 / AEI /10.13039/501100011033/ FEDER, UE, and the Department of Research and Universities of the Generalitat de Catalunya code 2021 SGR 01536.

