## Supplementary materials for "Clustering individuals using INMTD: a novel versatile multi-view embedding framework integrating omics and imaging data"

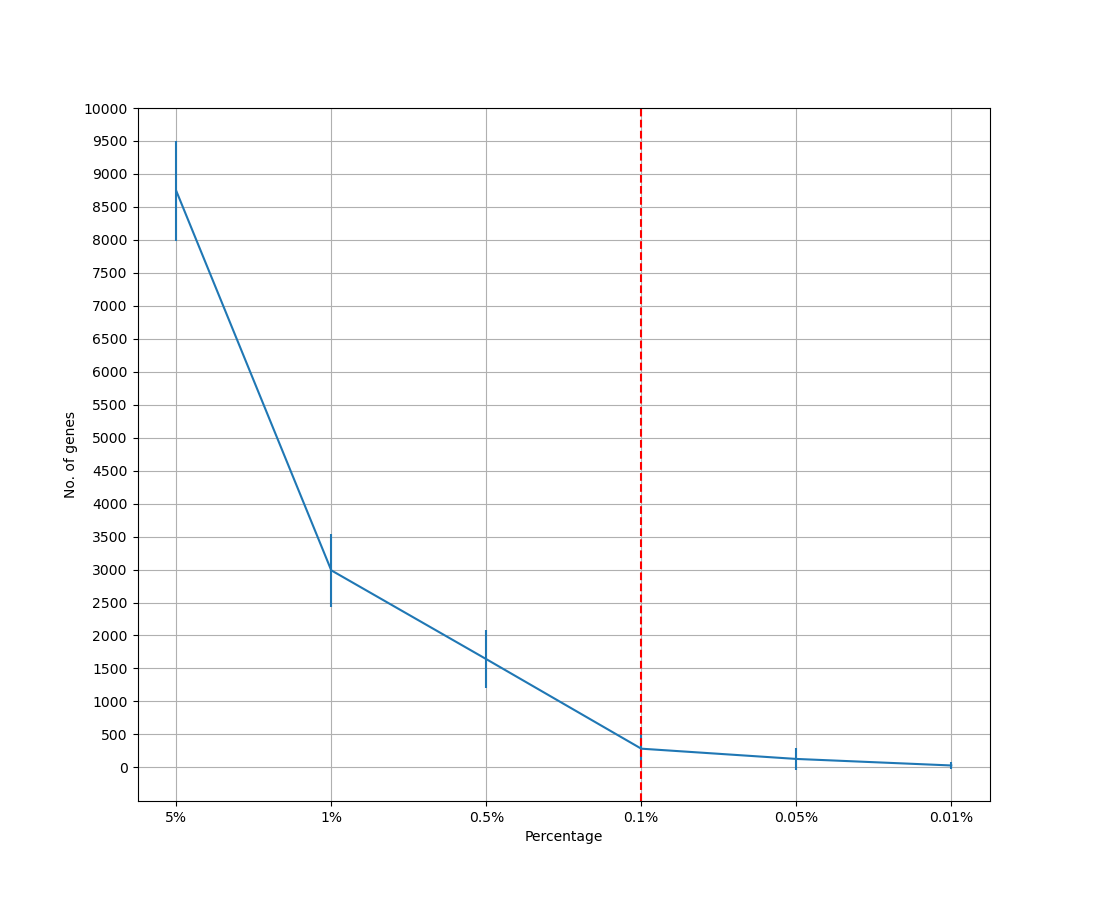

Supp. Fig. 1: Average No. of genes (y-axis) selected for each population subgroup with different thresholds (x-axis). The red vertical dashed line indicates the selected threshold.

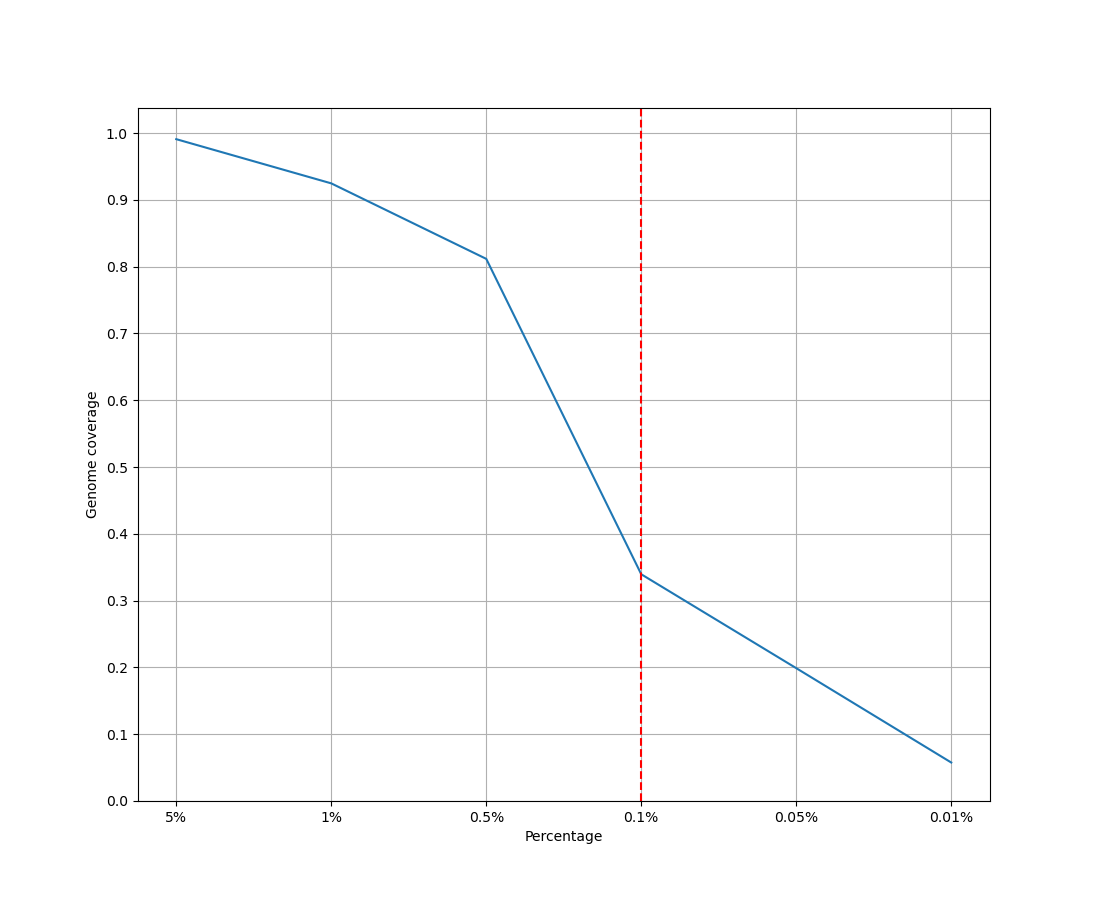

Supp. Fig. 2: Genome coverage of genes (y-axis) selected for all population subgroups with different thresholds (x-axis). The red vertical dashed line indicates the selected threshold.

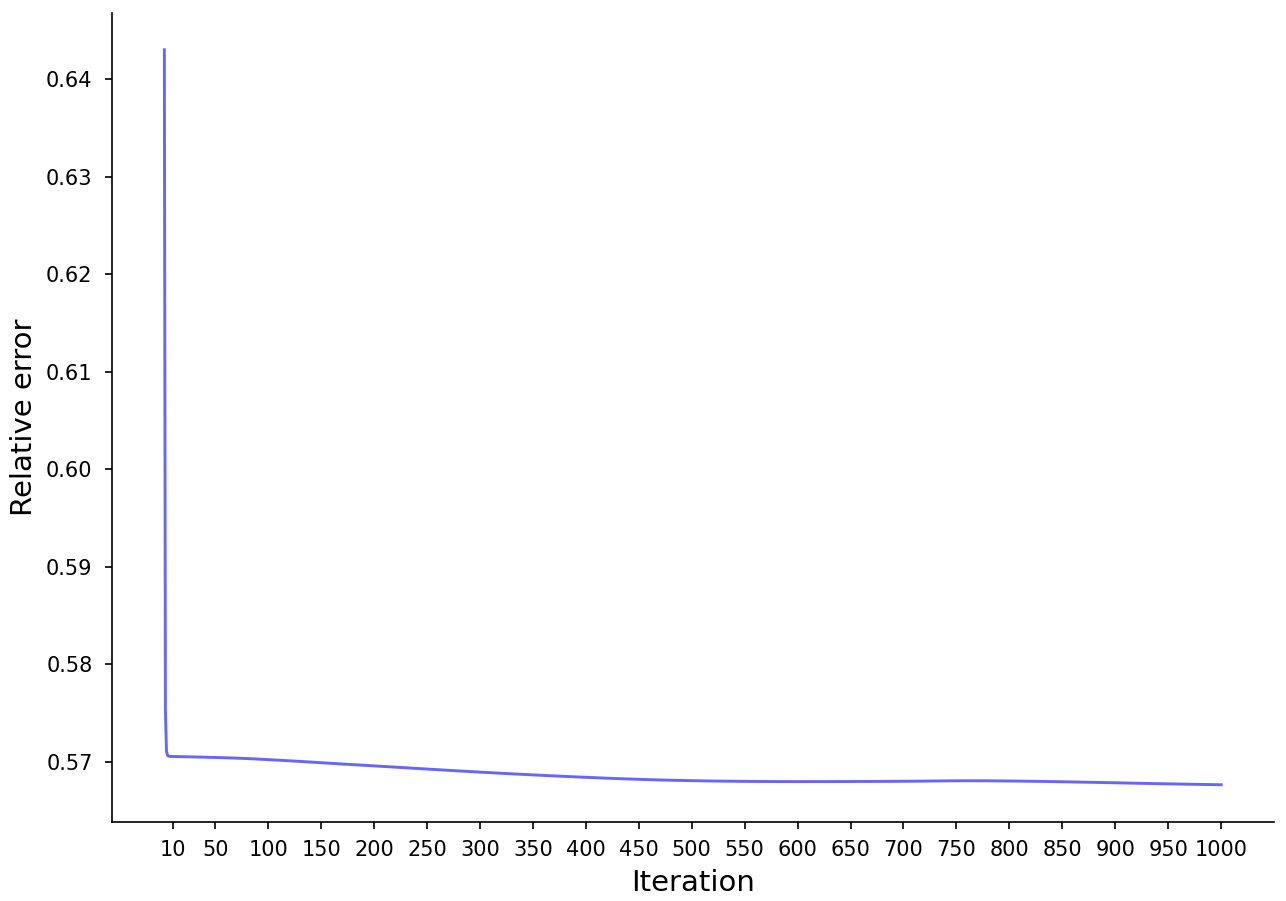

Supp. Fig. 3. Trajectory of the total relative error.

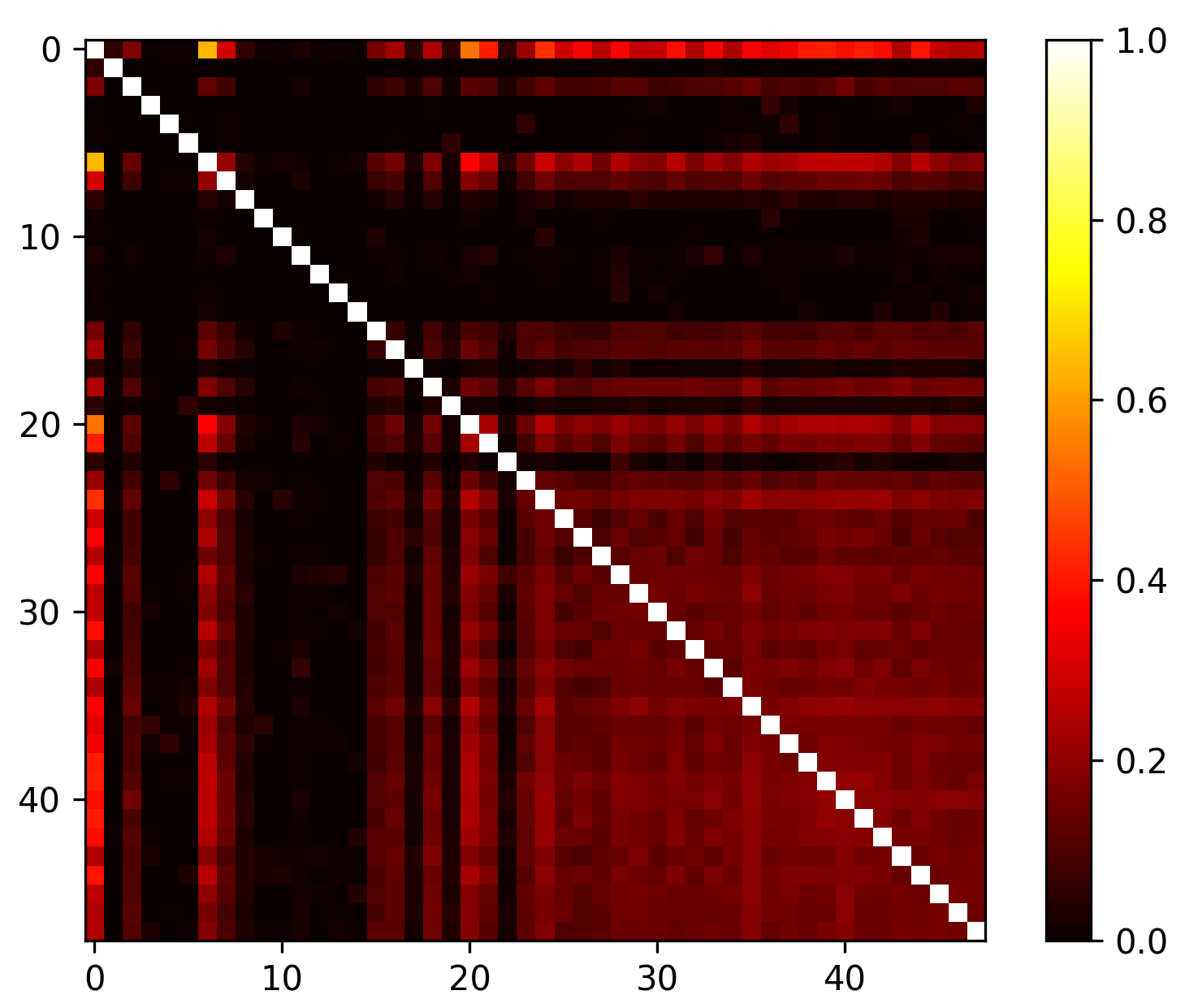

Supp. Fig. 4: Heatmap of the covariance matrix of $G_{1}$.

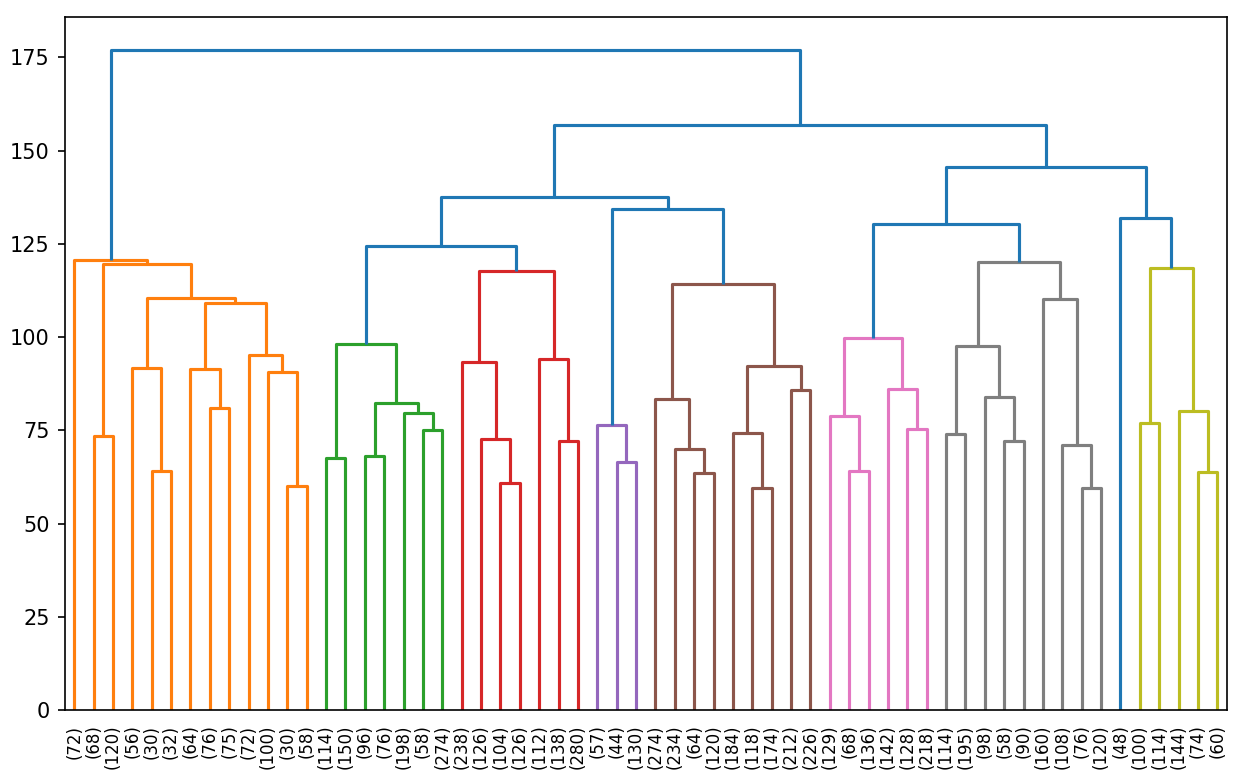

Supp. Fig. 5: The dendrogram of the hierarchical clustering based on $G_{3}$. Only the last 60 clusters formed in Ward’s linkage are shown and the size of every cluster is indicated in parentheses.

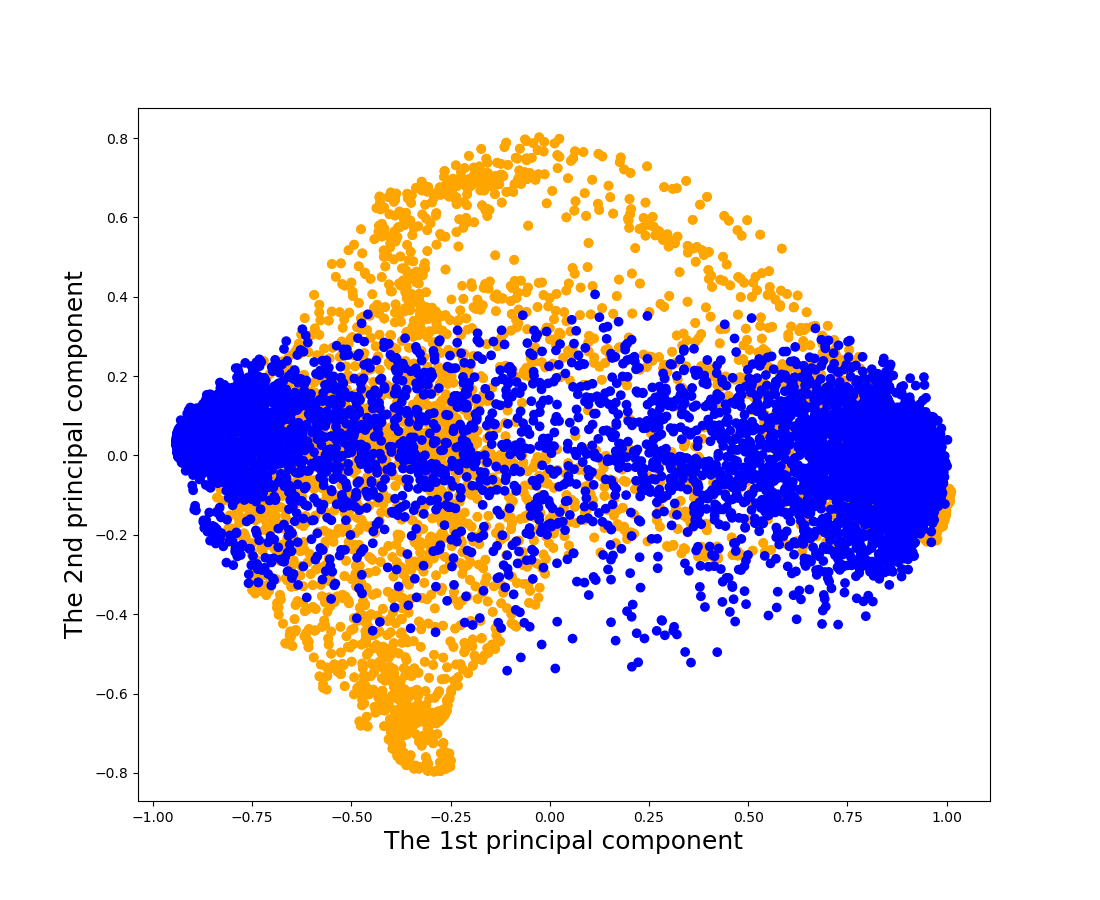

Supp. Fig. 6: The scatter plot of SNPs (in blue) and facial landmarks (in orange) with the first two principal components. The plot was created using principal component analysis (PCA) with cosine kernel on 7,160 randomly selected SNPs and all 7,160 facial landmarks in the space of $G_{1}$. The subsampling of SNPs is for balancing the two data types and the ease of visualization.

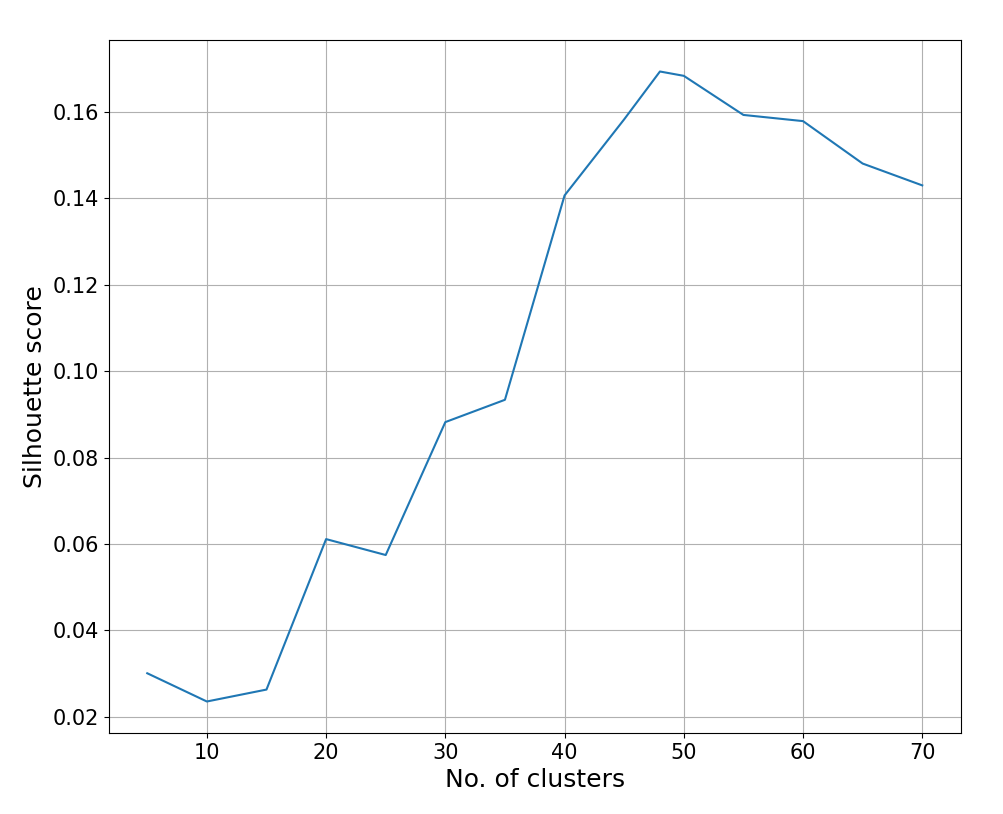

Supp. Fig. 7: Silhouette score (y-axis) for different number of clusters (x-axis) based on $k$-means clustering on $G_{1}$.

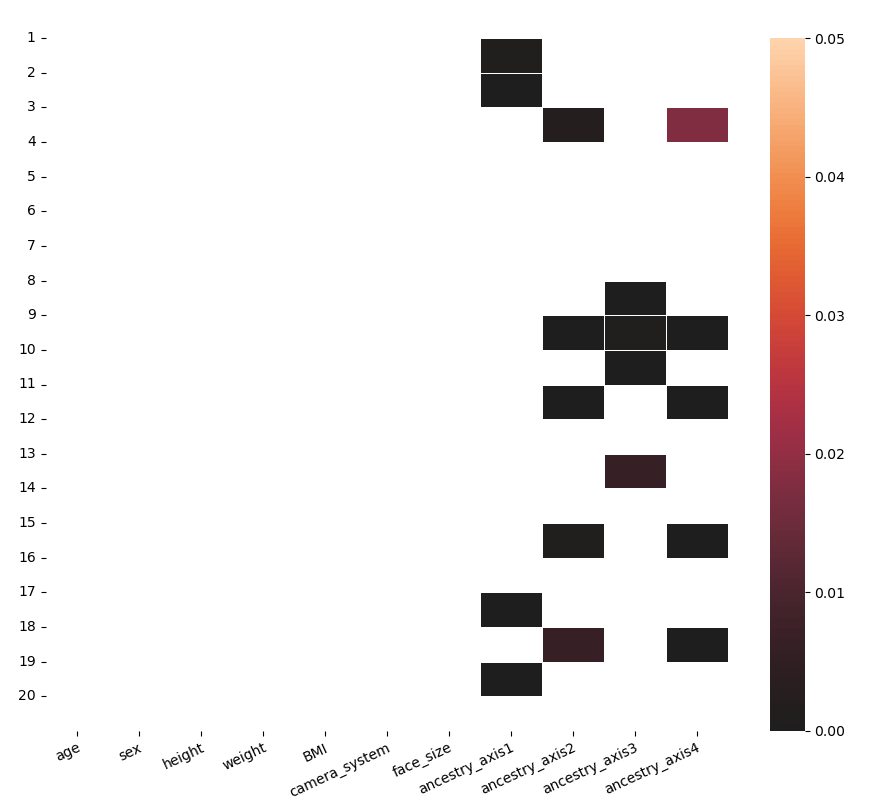

Supp. Fig. 8: Heatmap of P-values for the F test between every unconfounded $G_{1}$ embedding vector and every confounder and ancestry axis.

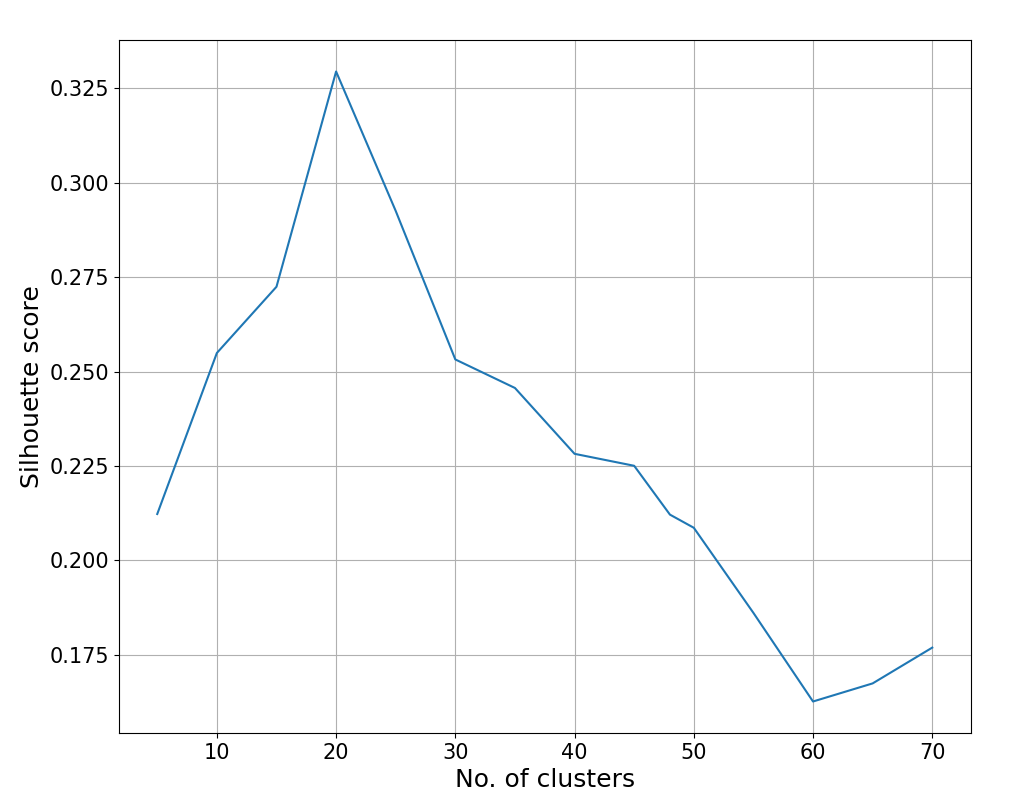

Supp. Fig. 9: Silhouette score (y-axis) for different number of clusters (x-axis) based on $k$-means clustering on the unconfounded $G_{1}$, which is composed of the 20 vectors of $G_{1}$ that are not significantly associated with any confounders.

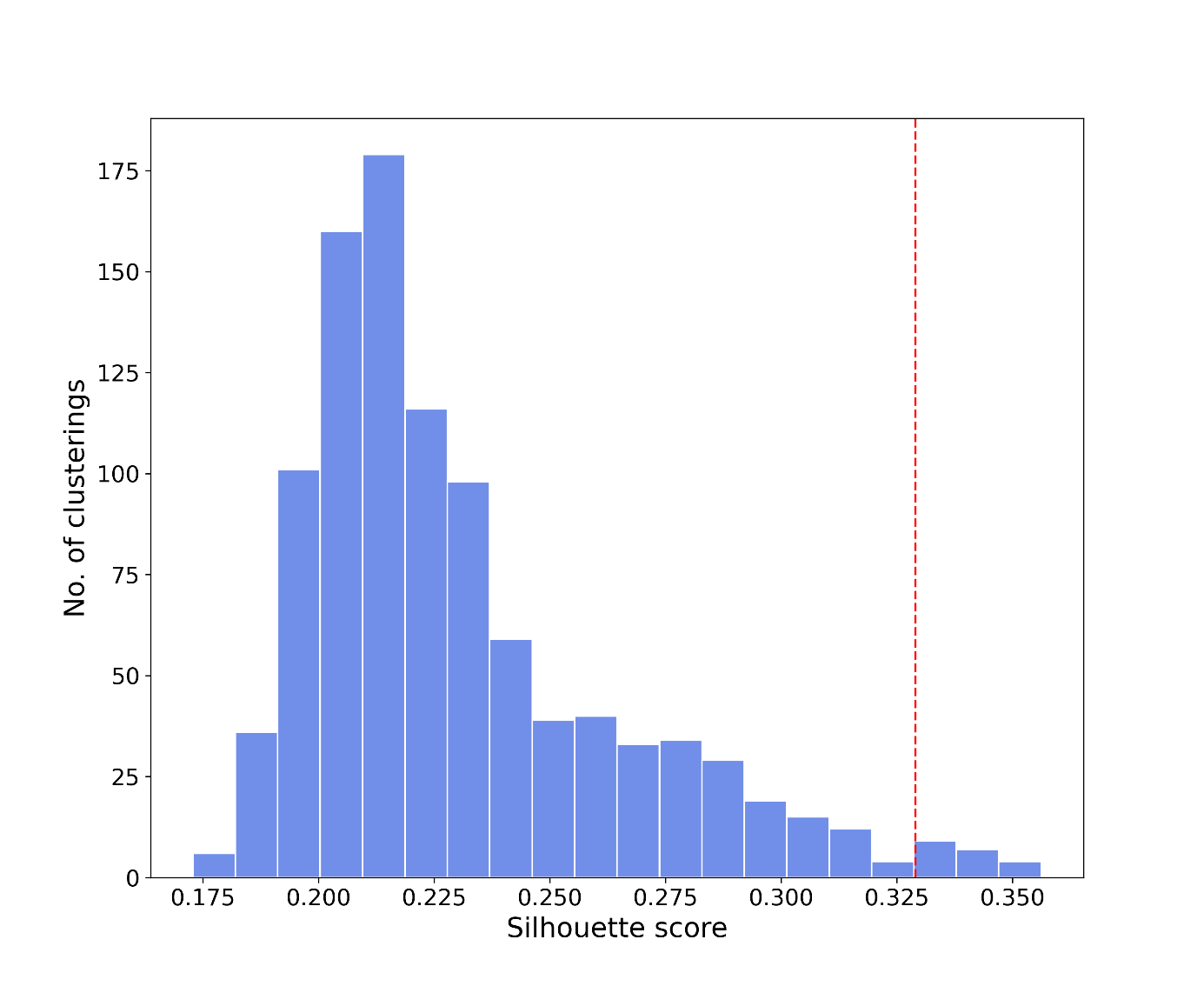

Supp. Fig. 10: Histogram of Silhouette scores from 1000 repetitions of $k$-means clustering based on 20 randomly sampled $G_{1}$ vectors. The number of clusters for each repetition is the same (20). The red vertical line indicates the Silhouette score (0.329) of the clustering derived from 20 unconfounded $G_{1}$ vectors, also with 20 clusters.

| **Subgroup No.** | **Adj. P-value** | **Subgroup No.** | **Adj. P-value** |
| --- | --- | --- | --- |
| 1 | **4.73e-04** | 11 | **1.27e-03** |
| 2 | **1.70e-08** | 12 | 1.00 |
| 3 | **7.81e-06** | 13 | 0.130 |
| 4 | **3.75e-03** | 14 | 1.00 |
| 5 | **2.60e-03** | 15 | 0.200 |
| 6 | **3.90e-02** | 16 | 5.83e-02 |
| 7 | **7.99e-08** | 17 | **4.77e-02** |
| 8 | 1.00 | 18 | 9.83e-02 |
| 9 | 1.00 | 19 | 0.638 |
| 10 | **3.75e-03** | 20 | **6.94e-07** |

Supp. Table 1: Adjusted log-likelihood ratio (LLR) P-values from logistic regression models for each population subgroup. For each subgroup, a binary variable (whether an individual belongs to this subgroup or not) was created as the dependent variable against four ancestry axes (plus intercept), modelled by logistic regression with the ‘lbfgs’ solver. All LLR P-values have been corrected for multiple testing via the Benjamini-Hochberg (BH) procedure. Adjusted P-values lower than 0.05 (threshold for significance) are in bold.
